# Profiling of bacterial and fungal microbial communities in cystic fibrosis sputum using RNA

**DOI:** 10.1101/336230

**Authors:** Nora Grahl, Emily L. Dolben, Laura M. Filkins, Alex W. Crocker, Sven D. Willger, Hilary G. Morrison, Mitchell L. Sogin, Alix Ashare, Alex H. Gifford, Nicholas J. Jacobs, Joseph D. Schwartzman, Deborah A. Hogan

## Abstract

Here, we report an approach to detect diverse bacterial and fungal taxa in complex samples by direct analysis of community RNA in one step using NanoString probe sets. We designed rRNA-targeting probe sets to detect forty two bacterial and fungal genera or species common in cystic fibrosis (CF) sputum, and demonstrated taxon-specificity of these probes as well as a linear response over more than three logs of input RNA. Culture-based analyses correlated qualitatively with relative abundance data on bacterial and fungal taxa obtained by NanoString and the analysis of serial samples demonstrated the use of this method to simultaneously detect bacteria and fungi and to detect microbes at low abundance without an amplification step. The relative abundances of bacterial taxa detected by analysis of RNA correlated with the relative abundances of the same taxa as measured by sequencing of the V4V5 region of the 16S rRNA gene amplified from community DNA from the same sample. We propose that this method may complement other methods designed to understand dynamic microbial communities, may provide information on bacteria and fungi in the same sample with a single assay, and, with further development, may provide quick and easily-interpreted diagnostic information on diverse bacteria and fungi at the genus or species level.

**Importance:** Here we demonstrate the use of an RNA-based analysis of specific taxa of interest, including bacteria and fungi, within microbial communities. This multiplex method may be useful as a means to identify samples with specific combinations of taxa and to gain information on how specific populations vary over time and space or in response to perturbation. A rapid means to measure bacterial and fungal populations may aid in the study of host response to changes in microbial communities.

## Introduction

Cystic Fibrosis (CF) is a life-limiting genetic disease associated with chronic lung infection that often leads to progressive lung function decline interspersed with pulmonary exacerbations (1-4). CF lung infections are often polymicrobial, heterogeneous between subjects, variable within individuals over time and can contain bacteria, fungi, and viruses (5-14). The characterization of microbes within CF sputum samples containing diverse bacteria and fungi has been performed using various methods including culturing on different media, sequencing of PCR amplicons for ribosomal RNA genes (rDNA) or ITS (internal transcribed spacer) sequences from bacteria and fungi, respectively, and metagenomic sequencing of community DNA. In this work, we assess the use of the NanoString nCounter technology (15) to detect specific bacterial and fungal taxa in sputum samples from individuals with CF-related respiratory infections.

The NanoString methodology detects RNAs using color-coded probe pairs that are optically detected. Together, a fluorescent reporter probe and a capture probe recognize an approximately one hundred base pair sequence within the target RNA. The target RNA-probe complexes are then captured and counted based on their unique fluorescent probe signatures (**Fig. 1**). Barczak *et al.* (16) previously demonstrated the use of this technology to detect species-specific mRNAs of bacteria, parasites, and fungal species and viruses in samples of blood and urine containing one pathogen, but not in clinical samples with complex communities such as CF sputum. In addition, NanoString-based methods were used to monitor seventy-five *Pseudomonas aeruginosa* virulence-associated mRNAs in RNA from CF sputum (17) and to detect pathogen mRNAs in samples from single-species mammalian infection models (18-21).

**Fig 1.**
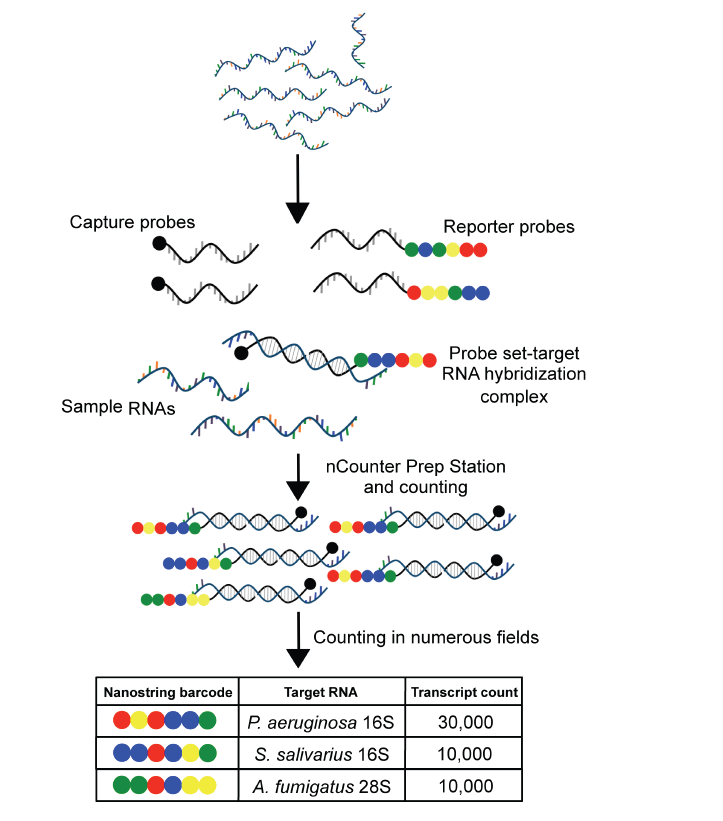
NanoString detection method schematic. Frozen samples are lyophilized and total RNA is extracted. Total RNA is mixed with the probe sets, with each containing a reporter and capture probe for each target RNA. After hybridization, the excess probes are removed and the probe-target complexes are aligned and immobilized in the nCounter cartridge. Fluorescent color sequences are counted and tabulated yielding counts that correspond to the number of target molecules.

We designed and tested taxon-specific NanoString probe sets for the simultaneous detection of forty-two CF-associated bacterial and fungal genera or species by monitoring levels of rRNAs. Probe fidelity was determined using control RNA mixtures as well as by comparison to results obtained by culture and previously established DNA-based community profiling methods. This technology can be used to monitor diverse taxa of interest, including but not limited to, bacterial and fungal targets, in complex microbe-containing samples. We demonstrate its use in monitoring changes in populations over time. Future applications of this approach include the development of improved diagnostics and strategies to monitor changes in microbial communities in response to perturbation.

## Results

### Probe selection and design

The bacteria and fungi commonly found in CF respiratory sputum have been well-described by culture and deep sequencing methods (5, 6, 14, 22-26). With this knowledge of species likely to be present, we designed a set of NanoString probes to detect taxon-specific rRNA sequences within total RNA isolated from sputum. In addition to the forty-two bacterial and fungal genera or species that are frequently detected in CF sputum, we included a few other taxa that commonly occur in non-CF respiratory samples (**Table 1**). In some cases, we designed genus-level probes (e.g. *Streptococcus* and *Staphylococcus*) in addition to probes that could detect specific species or complexes within these genera. For other taxa, such as *Burkholderia*, only a genus-level probe was included to streamline analysis and limit costs for this proof of concept study. For each taxon that we chose to monitor, a NanoString probe set was designed to target rRNA sequences from bacterial (16S or 23S rRNA) or fungal taxa (18S or 28S rRNA). Two probes that recognized adjacent sequences within a consecutive 100-nucleotide region were designed and one was modified to serve as the capture probe and the other modified to serve as the reporter probe (**Fig. 1**). The sequences for the two probes were designed to have >95% identity to the comparable region in all publically available genomes for the target taxon (inclusivity criteria) and less than 90% sequence identity to any non-ribosomal RNA sequence in the genomes of bacteria, fungi or humans available in Genbank at the time of probe design (exclusivity criteria) as described in the methods. The inclusivity and exclusivity assessments are presented in **Table S1**.

**Table 1.** Taxa detected with the NanoString probe sets

| **Bacteria** | **Fungi** |
| --- | --- |
| *Achromobacter xylosoxidans* | *Aspergillus fumigatus* |
| *Actinomyces odontolyticus* | *Candida albicans* |
| *Anaerococcus prevotii* | *Candida dubliniensis* |
| *Burkholderia* spp. | *Candida glabrata* |
| *Corynebacterium pseudodiphtheriticum* | *Candida lusitaniae* |
| *Fusobacterium nucleatum* | *Candida tropicalis* |
| *Gemella haemolysans* | *Candida parapsilosis* |
| *Gemella morbillorum* | *Exophiala dermatitidis* |
| *Haemophilus influenza* | *Malassezia globosa* |
| *Haemophilus parainfluenzae* | *Malassezia restricta* |
| *Mycobacterium* spp. | *Rhodotorula glutinis* |
| *Mycobacterium abscessus* | *Rhodotorula mucilaginosa* |
| *Neisseria* spp. | *Scedosporium apiospermum* |
| *Porphyromonas catoniae* | *Trichosporon mycotoxinivorans* |
| *Prevotella melaninogenica* |  |
| *Prevotella oris* |  |
| *Pseudomonas aeruginosa* |  |
| *Ralstonia pickettii* |  |
| *Rothia dentocariosa/mucilaginosa* |  |
| *Staphylococcus* spp. |  |
| *Staphylococcus aureus* |  |
| *Stenotrophomonas maltophilia* |  |
| *Streptococcus* spp. |  |
| *Streptococcus oralis/mitis/pneumonia* |  |
| *Streptococcus anginosus* |  |
| *Streptococcus intermedius/constellatus* |  |
| *Streptococcus salivarius* |  |
| *Veillonella atypica* and *Veillonella dispar* |  |

### Analysis of specificity of probes using mixes of RNA from single-species cultures

To accompany the computational analysis, which suggested that taxon-targeting rRNA probes would be specific based on exclusivity and inclusivity criteria, we performed control experiments using mixes of total RNA from single species cultures of twelve bacteria and two different fungi to assess potential cross-reactivity. We prepared mixtures of RNA extracted from single species cultures. Each mixture, containing equivalent amounts of RNA from three to five species, was analyzed using the panel of NanoString probe sets. The species RNA included in each mix are indicated in **Fig. 2** (see **Table S2** for strain information).

**Fig 2.**
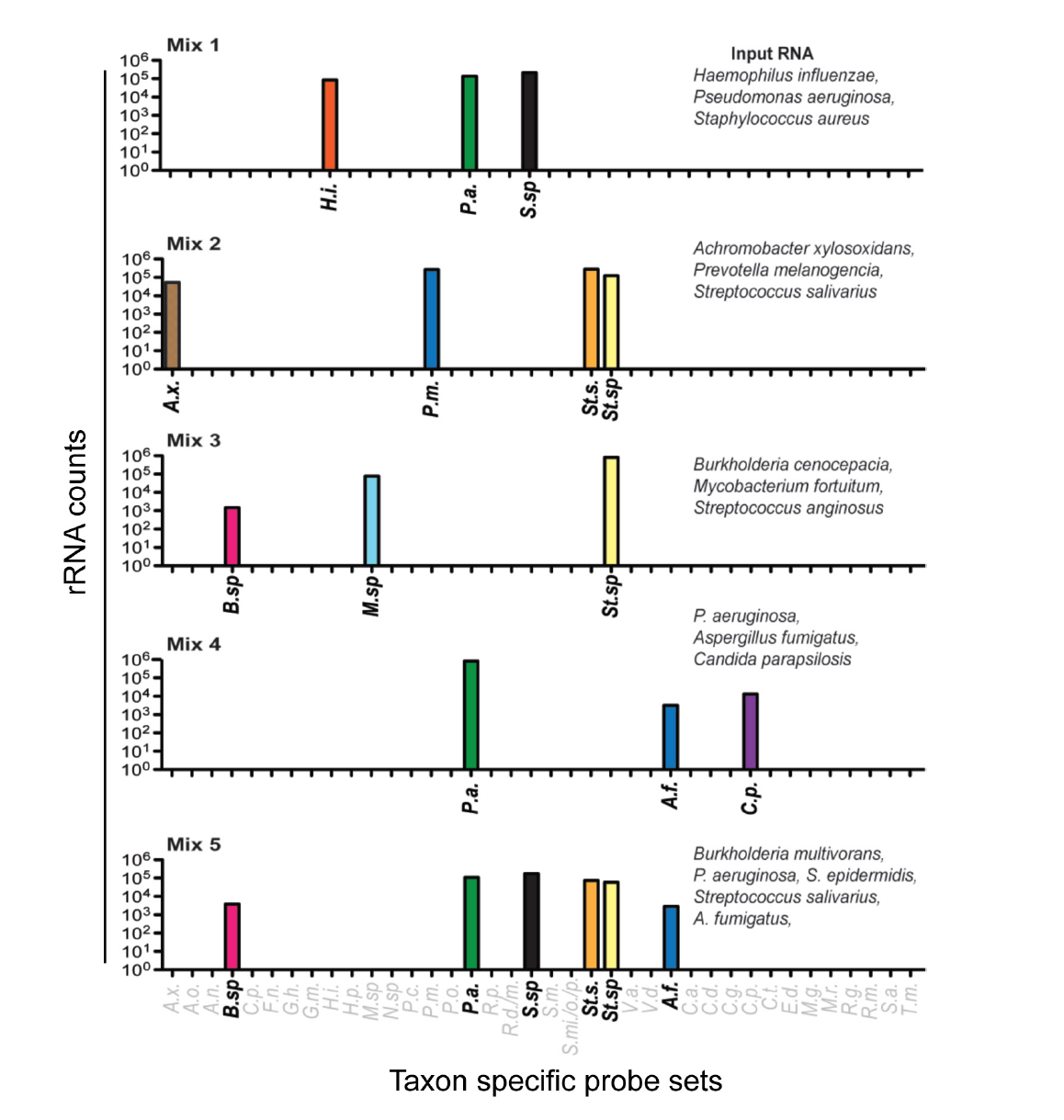
Specificity of select bacterial rRNA targeting probe sets. Control mixes were prepared with total RNA from three to five species per mix and analyzed using the full panel of probe sets. Strain details are in Table S2. The probe sets for taxa not present in any of the RNA mixes (in grey font) yielded no counts above background, indicating no cross-reaction with RNAs in the mixes. All data are presented as counts, and all taxon abbreviations are defined in Table S1. The species detected by each probe set is defined by first letters of the genus and species name, except for *Streptococcus*-targeting probe sets which are abbreviated “St.” to differentiate from *Staphylococcus*-targeting probe sets which are abbreviated “S”. Probe sets that detect multiple species within a genus are indicated by “sp.” as for the probe sets that detect *Staphyloccocus* species (*S.sp.*), *Streptococcus* species (*St.sp.*) and *Burkholderia* spp. (*B.sp.*).

Taxon-specific probe sets gave strong positive signals, measured as counts, when cognate rRNAs were present in the mix (**Fig. 2**). The average positive signal was 1000-fold higher than the signals from other probe sets. There were no cases of false positives due to cross-reaction with other probes (counts for all probes shown in **Table S3A**). As shown for Mixes 2 and 5, the *Streptococcus salivarius* RNA reacted with the species specific *Streptococcus salivarius* probe (St.s) and the *Streptococcus* spp. probe (St. sp.), while *Streptococcus anginosus* only reacted with the genus level *Streptococcus* spp. probe as no species-specific probe was present. Both *Staphylococcus aureus* and *Staphylococcus epidermidis* reacted with the *Staphylococcus* spp. (S.sp) probe. Two different *Burkholderia* species, *B. multivorans* and *B. cenocepacia*, were detected by the *Burkholderia* species probe (B. sp.). Each mixture that included *Pseudomonas aeruginosa* included RNA from a distinct clinical isolate (strains detailed in **Table S1**), and each reacted equally well with the species-specific *P. aeruginosa* probe. *Candida parapsilosis* and *Aspergillus fumigatus* RNAs reacted specifically with their cognate probes, and did not cross-react with probes from other *Candida* or *Aspergillus* species, respectively.

To assess the range of counts for which probe set signals were linear, we analyzed RNA Mix 1, which contained RNA from *Pseudomonas aeruginosa, Haemophilus influenzae*, and *Staphylococcus aureus*, with total RNA amounts from 0.0005 ng to 5 ng. We found that the probe sets for *P. aeruginosa, H. influenzae* and *S. aureus* all yielded the expected linear increase in counts that correlated with sample input with the linear range between 100 and >70,000 counts (R^2^>0.999 for all three taxa) (**Table S3A**).

### RNA-derived NanoString profiles were consistent across replicates

As a first step towards evaluating the NanoString panel above for the analysis of microbial communities within sputum, we obtained de-identified sputa that were expected to contain many of the taxa targeted by the probes that we had designed. We extracted total RNA from five sputum samples (sputum samples A-E), and analyzed the RNA in two independent runs. In every case, we found that rRNA counts for each bacterial and fungal taxon strongly correlated between replicates indicating the high reproducibility of the method (**Fig. S1A,** R^2^>0.999 each sample). The counts from bacteria-and fungus-targeting probes are plotted separately (**Fig. S1A**) as the counts for bacterial RNAs were 100-to 1000-fold higher than those for the fungi suggesting that the bacteria were more abundant. In addition, we separately recovered RNA from homogenized sputum that had been divided into three aliquots (samples F, G and H). Again, we found that the relative abundances of different taxa present in the three replicates from each sample were highly similar (**Fig. S1B,** R^2^>0.9 in each pairwise comparison per patient).

In the NanoString method, probes bind directly to their RNA targets without a cDNA synthesis step or any other enzymatic manipulation. Double-stranded DNA is not available for hybridization (**Fig. 1**). To confirm that the NanoString analysis of rRNA was detecting RNA and not DNA, three sputum-derived RNA samples were analyzed before and after RNase treatment. Total number of counts for all probe sets above background was reduced by >99% for all samples after RNase treatment and the community patterns determined by NanoString were markedly altered after RNase treatment indicating that RNA, and not DNA, was giving rise to the detected counts (**Fig. S2 and Table S3B**).

### Simultaneous analysis of bacteria and fungi using NanoString in sputum sample series from six patients hospitalized for disease exacerbations

One of the impetuses for designing this RNA-based method for complex culture analysis of CF sputum is to overcome several challenges associated with current culture based protocols including 1) the need to analyze CF samples using at least six different culture media (Blood agar, Chocolate Agar, MacConkey Agar, Saboraud Dextrose Agar, Mannitol Salt Agar, and OFPBL (Oxidation/Fermentation-Polymyxin-Bacitracin-Lactose) medium to detect potential pathogens of interest, 2) the inability to readily identify some pathogens, such as *Achromobacter xylosoxidans*, by growth characteristics alone, 3) the wide variation in growth kinetics and colony morphology types within a species, and 4) the use of a small, fixed volume of sputum which may limit the detection of less abundant or heterogenously-distributed species. To both assess the accuracy of NanoString 16S rRNA analysis as a technique for detecting species of interest within a mixed community and for assessing changes in sputum microbiota across samples from the same subject, we obtained and analyzed sample series from six hospitalized individuals undergoing treatment for disease exacerbation (designated E1 to E6). In each series, a sample was obtained on the first day of treatment then at least one more and as many as six more samples were obtained on subsequent days. A total of twenty-five sputa were analyzed. In addition to the clinical culture analysis of the Day 0 sample (**Table 2**), all specimens were plated on Sheep Blood Agar (SBA), Pseudomonas Isolation Agar (PIA) and Sabouraud Dextrose Agar (SDA) in our research lab (culture plates shown in **Fig. S3)**. In addition, total RNA was extracted from 100 µl of each sputum sample and 18 ng of total extracted sputum RNA was analyzed by NanoString. All probe sets were included in the analysis of each sample. **Fig. 3A** presents data for all positive signals.

**Table 2.** Clinical culture report for Day 0 samples from E-series. MSSA represents methicillin-sensitive *Staphylococcus aureus* and MRSA represents methicillin-resistant *Staphylococcus aureus*.

| Day 0 sample | Clinical culture report |
| --- | --- |
| E1 | Moderate *Exophiala dermatitidis* and few MSSA |
| E2 | *Achromobacter xylosoxidans* |
| E3 | *Pseudomonas aeruginosa* |
| E4 | *Burkholderia cenocepacia*, MSSA |
| E5 | MRSA, mucoid *Pseudomonas aeruginosa* |
| E6 | MRSA, MSSA, mucoid and non-mucoid *Pseudomonas aeruginosa* |

**Fig 3.**
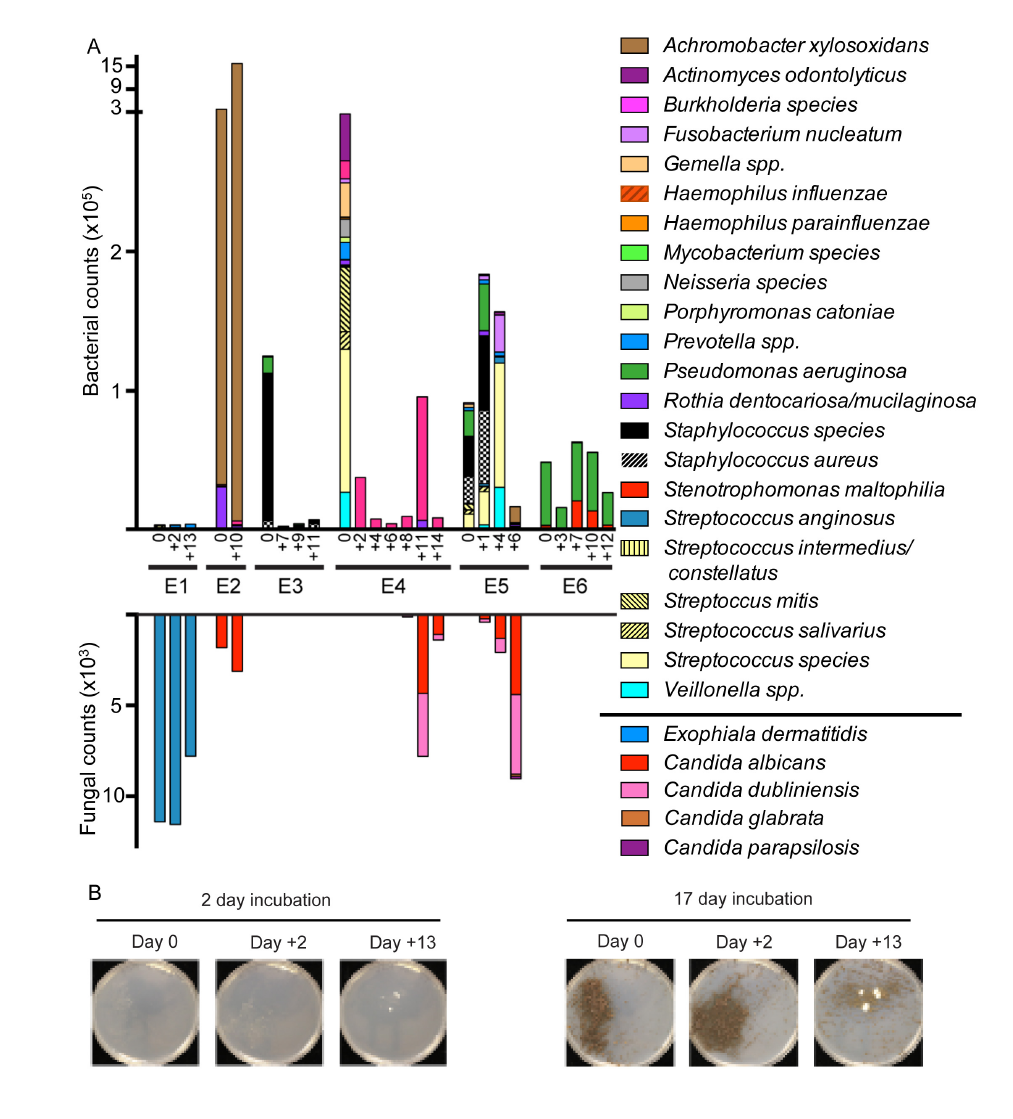
Bacterial and fungal community composition in exacerbation series from six subjects. **(A)** Serial sputum samples from six individuals with CF were obtained during treatment for pulmonary exacerbation and analyzed by NanoString (data in **Table S3C**). Microbes present in multiple subjects or abundantly in at least one subject are graphed as total counts normalized to positive controls. Bacteria and fungi are plotted on the upper and lower y-axes, respectively, though the data for bacterial and fungal probe sets were obtained in the same reaction. **(B)** Exacerbation series E1 showed high levels of *Exophiala dermatitidis* by plate culture only after extended incubation.

The analysis of sputum RNA from the E1 series showed high counts for the fungus *Exophilia dermatitidis* (*E.d.*) with no counts above background for any other fungi; the counts for *E. dermatitidis* were ∼100-fold higher than for any bacterial species suggesting that *E. dermatitidis* was a dominant microbe in these samples (**Fig. 3A**). While *E. dermatitidis* was not evident on culture plates after 48 hours of incubation at 37°C, the characteristic brown colonies of *E. dermatitidis* were clearly abundant on both SBA and SDA after more than a week (**Fig. 3B**).

*E. dermatitidis* was in the final clinical culture report of the Day 0 sample (**Table 2**).

In the two samples from the E2 series, NanoString counts for *Achromobacter xylosoxidans* were ten-fold higher than any other bacterial or fungal species at both time points (**Fig. 3**), and *Achromobacter xylosoxidans* was the only species detected in the clinical microbiology analysis of the Day 0 sample. Small, non-descript white colonies consistent with those formed by *Achromobacter xylosoxidans* were also detected on blood agar, PIA and SBA plates in our culture based studies (**Fig. S3-E2**). Low counts for *Candida albicans* were also detected and yeast colonies were evident on the SDA medium plates (**Fig. S3**).

In series E3, the most abundant NanoString signal from CF-associated pathogens was *P. aeruginosa* that are routinely reported clinically (**Fig. 3**-green bar); this species was the only species reported in the clinical microbiology analysis of the Day 0 sample and was clearly evident on the Day 0 sample on PIA (**Table 2** and **Fig. S3-E3**). *P. aeruginosa* was not detected by either NanoString or by culture at the later time points. The genus-level *Staphylococcus* probe (black bar) set gave very high counts and the *Staphylococcus aureus* (black hatched bar) probe set yielded ten-fold fewer counts in the Day 0 sample. The presence of non-aureus *Staphylococcus* species was confirmed by the presence of large numbers of non-hemolytic *Staphylococci* on the blood agar plate at the Day 0 time point, and the number of staphylococcal colonies on the blood agar plate decreased concomitant with the reduction in *Staphylococcus* spp. NanoString signal (**Fig. S3-E3**). Interestingly, the *S. aureus* NanoString counts remained very low but constant over the samples examined, suggesting that this signal was not due to cross reaction with other RNA from other *Staphylococcus* spp. that grew on the blood agar plates.

In samples from subject E4, counts for *Burkholderia* (pink bar) were high in the RNA from the sputum samples (**Fig. 3**); *Burkholderia* colonies that were slow-growing, characteristic of small colony variants, began to emerge on SBA after 48 h (**Fig. S3-E4**). The weak growth observed on plates from E4 was confirmed to be *Burkholderia* by 16S rRNA gene sequencing of amplified DNA from single isolates and the clinical microbiology reported *Burkholderia cenocepacia* as the dominant pathogen with “few” *S. aureus*. *Staphylococcus aureus* counts were elevated in the Day 0 sample, but not above the conservative threshold set in this study (see methods) (**Table S3C**). *Candida* colonies on SDA were more abundant at later time points and the signals from *Candida albicans* and *Candida dubliniensis* increased concomitantly in samples from later days. We confirmed that both *C. albicans* and *C. dubliniensis* were present by sequencing of the ITS1 region amplified from genomic DNA from individual colonies.

In samples from series E5, *Pseudomonas aeruginosa* (green bar), *Staphylococcus* spp. (black bar) and *Staphylococcus aureus* (black hatched bar) represented the strongest signals in the early time points (Day 0 and Day 1) (**Fig. 3-E5**). Both *P. aeruginosa* (mucoid) and *S. aureus* (MRSA) were detected in the clinical microbiology analysis of the Day 0 sample (**Table 2**). *P. aeurignosa* was detected on PIA medium and non-aureus staphylococci were evident on the SBA plate in our laboratory analyses (**Fig. S3-E5**). E5 sample from Day 6 showed *P. aeruginosa* and *S. aureus* signals to be 100-fold lower than on Days 0 and 1, and the counts for *Achromobacter* (*A.x.*-brown bar) were the highest among all taxa at this time point (>7000 counts) (**Fig. 3** and **Table S3C**). Small grey colonies that are typical for *Achromobacter* appeared on the PIA and SBA plates at these late time points (**Fig. S3**). *C. albicans* and *C. dubliniensis* were detected by NanoString with increasing levels at later days and this was concordant with the culture analysis (**Fig. S3-E5**). We confirmed that both of these species were present by sequencing of the ITS1 region amplified from DNA from individual colonies. *Candida glabrata* and *Candida parapsilosis* were detected, though at low levels, in E5 by both NanoString and their presence among the colonies on the Day 6 SDA plate was confirmed by sequencing of the ITS1 sequence amplified from the DNA of single isolates. The presence of multiple *Candida* spp. within single CF sputum samples is consistent with previous reports (14).

In series E6, the strongest signals were for *P. aeruginosa* (green bar) and *Stenotrophomonas maltophilia* (red bar) with much lower, but positive signals from *Achromobacter xylosoxidans* and *Burkholderia* species (**Fig. 3**). Of these taxa, only *P. aeruginosa* was reported in the clinical microbiology report for Day 0 (**Table 2**). The clinical lab reported isolation of *Staphylococcus aureus* from the E6 Day 0 culture (**Table 2**) and while the NanoString counts were above the highest background count, they were much lower than for the other bacterial pathogens (**Table S3C**). The levels of *S. maltophilia* rRNA increased on days 7 and 10. Bacterial growth from the E6 samples was slow and weak on Blood agar plates, and species determination of the colonies that finally emerged was not performed (**Fig. 3-E6**). *Candida* colonies were only observed on Days 0 and 3 and a positive NanoString signal for *Candida albicans* was elevated (**Fig. S3**) but below the threshold we had set.

Together, these data indicate that all of the pathogens reported upon clinical microbiology analysis of the Day 0 sputum samples were present and abundant in the NanoString data, with the exception of reports of *Staphylococcus aureus* in the clinical cultures from E1 and E6 (**Table 2**). *Staphylococcus aureus* was not evident in our laboratory culture of a separate aliquot of the same sample. The NanoString counts for the predominant bacterial pathogen were high (>3,800 counts) regardless of whether the strains grew robustly on medium, such as *P. aeruginosa* in E3, or slowly, such as *Burkholderia cenocepacia* in E4, *Pseudomonas aeruginosa* in E6 and *Exophiala* in E1. Together, these data indicate that the NanoString methodology can specifically detect bacteria (E2-E6) and fungi (E1, E2, E4, and E5) within samples using a single assay.

In addition, the NanoString approach can provide insights into dynamics of bacterial and fungal species abundance over time or in response to treatment, especially when one taxon is of lower abundance. As one example, in subjects E4 and E5, the number of *Candida* colonies and *Candida* NanoString counts increased over the course of treatment especially when sputum was analyzed on SDA plus gentamicin, a medium that suppresses the growth of bacteria (**Fig. S3**) consistent with previous reports of bacterial suppression of fungal growth in clinical cultures (27). The only fungi detected among these samples were *Exophiala* and *Candida* spp., and no samples had counts above background from probe sets for other fungi encountered in CF samples (*Aspergillus fumigatus, Scedosporium* and *Trichosporon*). Except for samples from the E1 series, the total probe counts from all fungal rRNA-specific probe sets were more than 1,000x lower than the total counts from probe sets designed to target bacterial rRNAs.

### Relative abundance of CF pathogens determined by NanoString correlates with Illumina sequencing of community DNA and culture-based analysis of sputum

Eighteen sputum samples collected during outpatient visits were divided for separate RNA and DNA extractions. We compared data obtained by NanoString-analysis of RNA to data obtained by well-validated deep sequencing analysis of 16S rRNA sequences amplified from DNA. Comparison to clinical microbiological culture data was also performed. We only focused on the bacteria in these analyses as the fungi were predicted to be of low abundance in most samples based on the clinical culture analysis. Total RNA was analyzed using the NanoString probes described above and the total DNA was subjected to amplification and sequencing of V4-V5 16S rRNA-encoding genes by Illumina HiSeq (**Fig. 4A**). Clinical culture analysis of the eighteen samples found that most of the bacterial pathogens frequently reported from CF sputum cultures were detected in at least one sample including *P. aeruginosa, Achromobacter xylosoxidans, Burkholderia* spp., *Ralstonia* spp., *Stenotrophomonas maltophilia,* and *Staphylococcus aureus*; two other commonly-reported CF pathogens were not detected *Haemophilus influenzae* and *Mycobacterium* spp. in the clinical microbiology analyses.

**Fig. 4.**
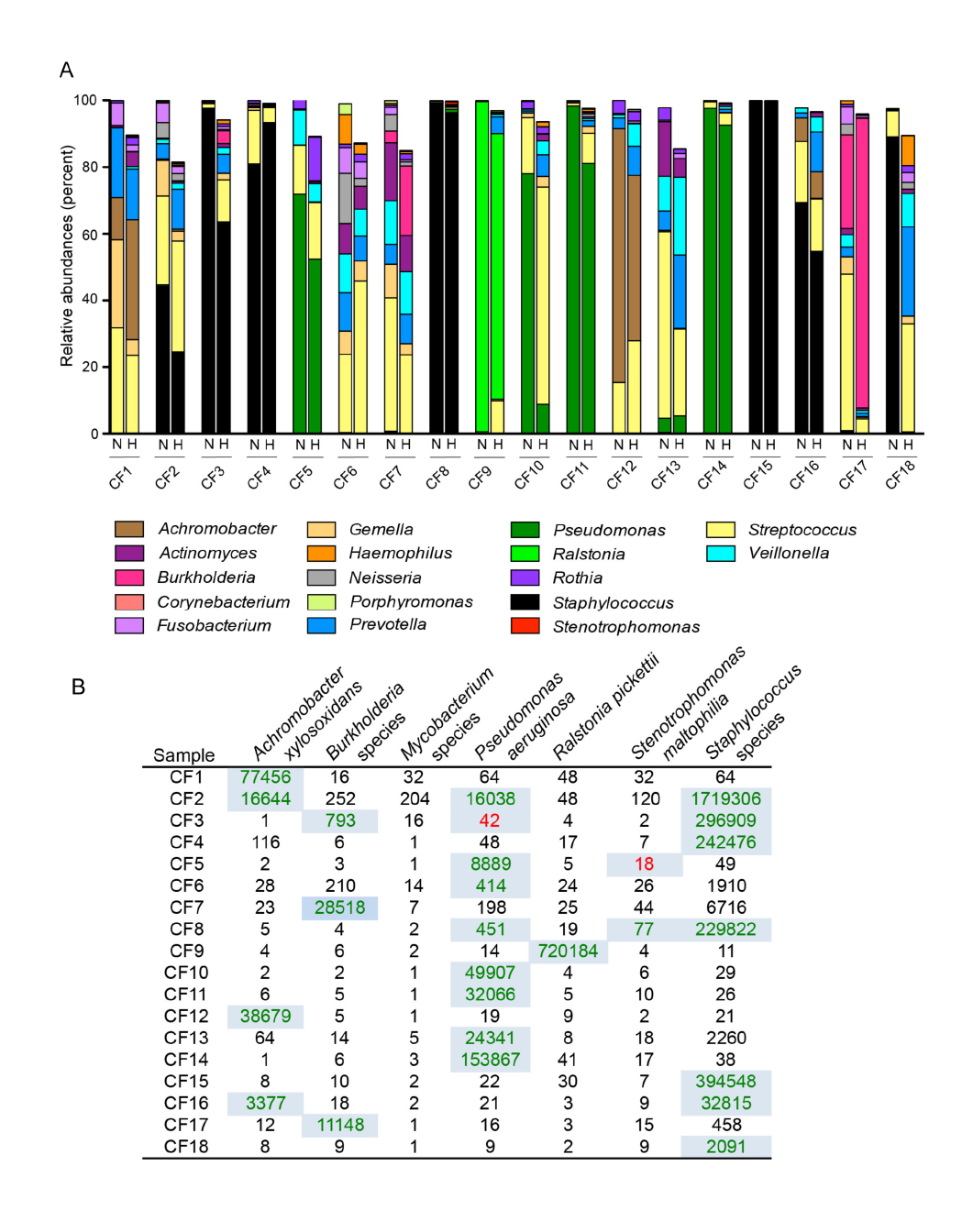
Bacterial community profiles detected by NanoString and 16S rDNA deep sequencing (HiSeq) methods correlate. **(A)** Eighteen CF sputum samples, labeled with the subject number, were analyzed by both NanoString (N) and 16S rDNA deep sequencing (HiSeq, H). Percent relative abundance of taxa detected by both methods is shown. Only the taxa measured in our NanoString code set are included in the deep sequencing profiles, but relative abundance was determined as a percentage of total reads, not only reads represented in the NanoString code set (data in Table S4). In cases where NanoString probes detected different species within the same genus, counts from individual species-specific probes were summed to correspond with the genus-level assignments for the HiSeq data. **(B)** The counts for probe sets correlating with CF pathogens are shown. Highlighted numbers indicate that the corresponding pathogen was also detected by clinical culture analysis. The numbers in red in CF3 and CF5 were reported as being “rare” in the clinical microbiology analysis and were at very low levels (<0.5%) in the HiSeq-based community profiles for those samples.

All of the genera at greater than 5% abundance in the Illumina-based community profiles of CF sputum were the taxa monitored by NanoString probes (**Table S4**). This result was not surprising since the taxa included in the NanoString codeset were selected based on their representation in prior Illumina-based community profiling of CF sputum. On average 92% ± 6% of the deep sequencing reads corresponded to taxa that were represented among the NanoString probes (**Table S4A**). For both data sets, the relative abundances were determined by division of the counts, for NanoString, or reads, for HiSeq, for the taxon of interest by the total number of counts or reads. For the HiSeq relative abundances, total reads, including those for taxa that were not assessed by NanoString, were used in the percent abundance calculation. Overall, there was high concordance between the data obtained by the two methods with an average Pearson correlation coefficient of 0.803 across all eighteen samples with all taxa, including both CF-related pathogens and species that are also frequent members of the oral microbiome. Together, these data indicate that there is minimal cross reaction between different NanoString probes. Furthermore, these data show that for CF sputum collected during outpatient visits, NanoString analysis of RNA and Illumina analysis of sequences amplified from DNA yielded similar results. Future studies will determine if data obtained from community RNA versus DNA differs in other contexts, such as over the course of intensive antibiotic treatment that might increase the fraction of non-viable or dead cells.

Clinical culture results from analysis of these eighteen samples described twenty-six instances in which one of the above bacterial pathogens was detected by culture and several samples contained two or three pathogens. For twenty-four of the twenty six bacterial pathogen detection events, there was a clear positive signal in the NanoString data that indicated the presence of the cultured pathogen (highlighted in green font with blue background in **Fig. 4B**); there was also a signal for the pathogen in the HiSeq data in each case. In two instances, culture analysis detected only “rare” colonies of *P. aeruginosa* (CF3) or *Stenotrophomonas maltophilia* (CF5), which can mean as few as one colony, and the NanoString signal was not clearly positive (red text with blue background in **Fig. 4B**). HiSeq analysis found *Pseudomonas* or *Stenotrophomonas* to be at very low relative abundance (0.01 and 0.2%) in samples CF3 and CF5, respectively. In one case (CF18), the pathogen in the clinical culture report was *Staphylococcus aureus* and this species was robustly detected by NanoString but at very low levels in the High Seq data (**Fig. 4A**). The reason for this discrepancy is not known, but it is worth noting that the sequnencing depth was not aberrantly low for this sample. These data suggest that nucleic acid based-methods, which involve the analysis of larger sample aliquots, may be capable of providing quantitative-or semi-quantitative information on the levels or activity of a taxon and may reinforce clinical microbiology data when few colonies of a species are observed in culture. One benefit of the NanoString method is that no amplification is performed prior to analysis aiding in quantitation. The high concordance between the NanoString, Hi-Seq, and culture data further support the validity of the probes and their use in describing microbial communities.

## Discussion

Here, we report a RNA-based method for detecting target species within complex microbial communities. We describe this approach for the analysis of sample types that have been well characterized by non-targeted approaches such as the sequencing of the metagenome or of amplified rRNA and ITS regions from community DNA in order to identify the taxa present. While the NanoString technology has been applied to the detection of mRNAs from microbial species within clinical samples or in tissues from animal experiments, we show the application of this technology to the analysis of rRNAs which are highly abundant molecules with sequences that are highly conserved across the genus and species level. In contrast to amplification based methods in which conserved primer-binding sequences are required, the regions detected within the rRNAs can be different for different taxa thus allowing for different taxonomic levels to be detected in the same multiplex reaction. Because of the use of a large (∼100 nt) region for hybridization to the two probes, computational analysis of the probes suggests that this method can accommodate some sequence divergence within species or genus, but this was not explicitly tested in these experiments.

Because fungi can grow slowly in laboratory cultures, as shown for the *Exophiala* in samples from subject E1, and may be unsuspected in some chronic contexts, a method that detects both bacteria and fungi has significant clinical value. We found that taxon-specific NanoString probe sets successfully detected fungi including *C. albicans, C. dubliniensis, C. glabrata, A. fumigatus*, and *Exophiala*. In our initial analyses using the NanoString method, we included probes that were complementary to ITS1 RNA sequences in addition to probes that bound fungal 28S rRNA (Table S1 for sequences). The ITS1 RNA counts were over 100-fold less abundant suggesting that ITS1 RNA, which is processed from rRNA containing RNAs, is less stable and thus less abundant than rRNAs.

We demonstrate significant concordance between this method and Illumina-based sequencing of amplified 16S rRNA sequences from community DNA for many of the organisms examined (**Fig. 4A**) including *Pseudomonas, Achromobacter, Burkholderia, Ralstonia, Stenotrophomonas, Staphylococcus, Actinomyces, Fusobacterium, Gemella, Neisseria, Rothia, Porphyromonas* and *Veillonella*. Consistent with a recent study that showed a relationship between deep sequencing and detection by culture when extended culture techniques were used (12), we found correlations between NanoString data and clinical microbiology culture assessment as well.

This method is most useful in the analysis of sample sets that have already been well-characterized by non-targeted approaches such as the sequencing of the metagenome or of amplified rRNA and ITS regions from community DNA. Once microbial community members of interest have been identified within a particular habitat, the number of tools available for gaining additional data on specific taxa within the community increases to include quantitative PCR, fluorescent in situ hybridization methods, and plate-based studies, among other methods. We propose that the NanoString method be added to the list of techniques for the analysis of specific taxa within a community and that it may be particularly useful for analyzing the relationships between microbes from different domains, which cannot be profiled using single primer sets, and between species within a genus which may lack resolution by other methods. Multiple probes for each taxon that target both small and large subunit rRNAs may enhance confidence in the detection of low abundance species.

In addition to the rRNA analyses described here, we have published a NanoString analysis of various *P. aeruginosa* mRNAs in clinical samples, and the inclusion of species-or strain-specific mRNAs could be useful in analyzing phylogenetically close taxa (17). The combination of probes for rRNAs and mRNAs, however, could be challenged by the difficulties in detecting lower abundance mRNA signals when rRNAs, which are orders of magnitude more abundant. Interfering complementary RNAs may be useful in reducing signals from highly abundant RNAs in order to enable the simultaneous analysis of probes for RNAs that are present in vastly different amounts, but we have not explored this strategy as part of this work. With further development, it is also possible to monitor host and microbial transcripts at the same time to better understand the interplay between host and microbe, particularly during changes in health.

The agreement between NanoString and 16S rDNA deep sequencing data for most taxa was striking considering the differences in target molecule (rRNA versus rRNA-encoding DNA) and the technological differences between the methodologies. A number of environmental community profiling studies have compared results from the analysis of extracted rRNA and rDNA sequences often with the goal of determining if RNA can be used to identify the more metabolically active members of the community (28-31). Ribosomal RNA, while more stable than mRNA, is degraded by stable, active RNases and thus it is tempting to speculate that the presence of RNA may serve as a more sensitive indicator of an active cell than the presence of DNA (32). If this is the case, the high concordance between NanoString and HiSeq data would suggest that the majority of microbes in the CF sputum samples from these stable patients are alive. While our data do not address whether RNA and DNA equivalently reflect the presence of live and dead cells, with taxon-specific probes in hand, studies that test this idea can now be performed.

This detection method, like any nucleic acid analysis method, has the potential for methodological bias. Complicating factors for the NanoString technology could include variable lysis, differences in transcription frequency of rRNA genes in different taxa, and differences in probe hybridization efficiency, thus validation strategies must be employed. While the bead beating based methods used here have been shown to be efficient at lysing fungi and other difficult-to-lyse taxa, the nucleic acid extraction methods used need to be assessed for other taxa that were not present in our sample sets such as *Mycobacteria*.

The studies described here represent the first step in employing the NanoString technology to the characterization of complex microbial communities using rRNA. In addition to analyzing and monitoring bacteria and fungi, host markers could also be added to the code sets to gain knowledge about the host state and response (33-36). Furthermore, the NanoString technology has already been introduced into clinical practice for the purpose of cancer cell profiling. While many additional validation experiments would need to be performed before a clinical diagnostic tool could be developed, the quick sample turnaround time and semi-quantitative outputs may allow for more rapid assessment of samples (as compared to traditional culture techniques, which may take days to weeks) as well as protocols for the assessment of the efficacy of treatments. This method provides direct quantitation of rRNAs present in a sample and thus can give insight into the number of rRNA molecules by sample weight, volume, or total RNA. Future studies are necessary to determine if rRNA levels correlate with meaningful metrics such as cell number or population activity.

One application of this methodology is the identification of community RNAs that contain significant levels of specific mixed-species microbial populations for use in more extensive analyses such as total RNA sequencing. In addition, the development of a sensitive methodology for the detection of fungi in human-derived samples can be readily adapted to address other research questions such as those regarding the roles of fungi in ulcerative colitis or asthma and reactive airway disease (37-41) as well as in environmental settings. Lastly, the development of taxon-specific probes that detect microbial rRNAs sets the stage for future studies aimed at understanding whether an analysis of microbial rRNA can provide an indication of the metabolic activity or pathogenic potential of specific microbial populations within complex communities.

## Materials and Methods

### Probe design

We detected microbial rRNAs using custom-designed species-and genus-specific probe sets. The probe sequences were designed by the NanoString probe design team to recognize all available sequences in Genbank for the taxon of interest and to not cross hybridize to off-target sequences in other bacteria, other fungi, or human genomes. All probe set sequences are listed in **Table S1**. Measurements were made using the NanoString nCounter system (NanoString Technologies, Seattle, WA) as previously described (15). As described in **Table S1**, the probes were purchased in batches, referred to as code sets and the code set composition was slightly modified over time to include additional taxa or to achieve different levels of specificity (for example replacing a genus level probe with a species level probe). Each probe contained either capture and detection elements. Each code set also contained six synthetic positive control probes that detect spiked transcripts added at a range of different concentrations and eight negative control probes that are not predicted to hybridize to any transcripts in the bacterial, fungal or human transcriptomes analyzed (NanoString Technologies).

### Analysis of RNA from in vitro-grown cultures

To assess the specificity of the RNA probes for specific species, equal amounts of purified total RNA from each microorganism was mixed, and 5 ng of the RNA mixture was used). The strains used in this study are listed in **Table S2**. RNA from laboratory cultures was prepared from cells harvested from 1-3 ml of liquid culture for all strains. *Pseudomonas aeruginosa* was grown in lysogeny broth (LB) with aeration on a roller drum. *Achromobacter xylosoxidans, Staphylococcus aureus, Burkholderia cenocepacia,* and *Mycobacterium fortuitum* were grown in tryptic soy broth (TSB) with aeration on a roller drum. *Prevotella melaninogenica* was grown under anoxic conditions using a GasPak EZ anaerobic container system and *Streptococcus salivarius,* and *Streptococcus anginosus* were grown statically in an atmosphere with 5% CO_2_ in Todd Hewitt broth amended with 0.5% yeast extract. *Haemophilus influenzae* was grown in Brain Heart Infusion (BHI) broth supplemented with 10 μg/ml NADH and 10 μg/ml hemin with static incubation in a 5% CO_2_ atmosphere. All cultures were inoculated from a single colony into 5 ml of the specified medium. Liquid cultures were grown at 37°C for 16-18 h except for *M. fortuitum* which was grown for 48 h. Cells were frozen and lyophilized prior to RNA extraction using the methods described in the following section.

### RNA and DNA isolation from sputum

Frozen aliquots (100 µl) were lyophilized for 5-16 h, as necessary. Lyophilized cells were disrupted using a mixture of 0.1 mm, 0.5 mm, and 1 mm zirconia/silicon beads (70 µl each) that was added to the dry pellet prior to agitation. Samples were lysed using 5 cycles of 30 sec, with 20 sec pauses in between, on an Omni Bead Ruptor Homogenizer (Omni International Inc.) set at 5.65 m/s. The disrupted sample was resuspended in Lysis Buffer (200 μl of 0.25 µg/µl lysostaphin and 3 µg/µl lysozyme in 10 mM Tris and 1 mM EDTA (TE) buffer, pH 8.0), and incubated at room temperature for 5-10 min. RNA isolation was then performed using Direct-zol kit (Zymo Research), with TRIzol, according to the manufacturer’s suggested protocol. RNA was eluted in two 20 µl volumes of elution buffer before freezing at -80°C until use. No DNAase treatment was performed. For RNA isolation reproducibility experiments, sputa were split into triplicate samples after homogenization, and then processed in parallel following the RNA isolation protocol described above.

For DNA isolation from sputum, lyophilized and disrupted sputum aliquots were resuspended in Lysis Buffer (described above) and incubated at room temperature for 5-10 min. Total DNA was isolated using Qiagen genomic DNA purification reagents, following the manufacturer’s protocol for Gram-negative bacterial DNA isolation. DNA pellets were resuspended in 40 μl TE buffer and frozen at -80°C until use.

### rRNA analysis using NanoString probe sets

We empirically determined that 15-20 ng total RNA was best for balancing the ability to detect less abundant species without overloading the optical detection method. For control mixes of RNA isolated from pure culture, 5 ng total RNA was analyzed, unless otherwise specified. Sample RNA was hybridized to the probe sets for 12-16 h at 65°C using the NanoString nCounter Prep Station instrument according to manufacturer’s instructions. Targets were counted on the nCounter using 255 fields of view per sample and output results were extracted using nSolver Analysis Software (NanoString Technologies).

### Background determination and detection of positive samples

A series of internal negative control probes that do not target sequences in any known organisms are included in every sample, and the signals associated with these probes inform the relative quality and background hybridization in each individual analysis. Following the manufacturer’s recommendation of using limits of detection based on the maximum signal for internal negative control probes, we chose a background subtraction method in which three times the maximum detection by internal negative controls (3xMax) was subtracted from all probe counts. Prior to analysis, a background subtraction of the 3xMax value was performed for all control mixes and patient samples. For comparisons of levels across samples, we normalized samples based on the mean of the positive controls.

### Patient cohort for CF sputum collection

The samples were collected in accordance with protocols approved by Center for the Protection of Human Subjects at Dartmouth. Expectorated sputum samples used in the comparison of NanoString and Hi-Seq methods were collected from adult subjects with CF during a routine office visit. Enrolled subjects provided two separate sputum samples during a single visit, one of which was analyzed by standard culture methods for this specimen type at the Dartmouth Hitchcock Medical Center clinical microbiology laboratory. The second sputum sample was divided into aliquots (one for DNA and one for RNA extraction) then frozen on-site using a portable -80°C freezer shuttle (Sterling Ultracold) prior to storage in an upright -80°C freezer. Sputum samples were stored at -80°C for at least 1 h and often for multiple days or weeks before analysis.

In serial samples collected from the same patient, samples were collected upon admission for treatment of a disease exacerbation as well as over the course of treatment. Sputum samples were frozen upon collection and stored at -80°C until samples were processed for RNA isolation as described below.

### DNA deep sequence analysis of bacterial 16S rDNA

Total genomic DNA (gDNA) isolated from 18 sputum samples was used as template for the amplification and analysis of the V4V5 regions of 16S rRNA encoding genes by deep sequencing at the Marine Biological Laboratory (Woods Hole, MA). Sequencing was performed on the HiSeq System (Illumina Inc., San Diego, CA) as previously described (42). Sequencing reads were processed and taxonomic assignments were called using a custom bioinformatic pipeline at the Marine Biological Laboratory. Briefly, raw reads were quality filtered as previously described (43, 44). Chimeric reads were identified and removed using the UChime (45) algorithm and *de novo* reference database modes. Taxon assignments were performed with GAST (43) against a curated SILVA database (46). Operational taxonomic unit (OTU) clustering was determined using UCLUST with a 97% similarity threshold (47). For the comparative analyses with the NanoString methods, a threshold of 0.01% was set for positive signals.

### Analysis of RNase treated RNA

To determine specificity of NanoString detection for RNA versus DNA, RNAse treatment was performed on 3 samples after RNA isolation (F, G, and H). For these three samples, 1 μl RNaseA was added to 5 μl of isolated RNA and incubated at 37°C for 1 h. The samples were then diluted 1:10 in 45 µl of 10 mM Tris-HCl (pH 7.5), 300 mM NaCl, 5 mM EDTA (pH 7.5) and 2 μl of RNase A/T1 mix (Thermo Fisher Cat # EN0551). This mixture was incubated for 1 h at 37°C. Non-treated RNA samples were then diluted 1:10 and equal volumes of RNAse treated and non-treated matched samples were used for the NanoString code set.

### Culture analysis of serial sputum samples

Approximately 10 µl of sputum was removed with metal Scienceware Microspoon and plated on Tryptic Soy Agar (TSA) with 5% Sheep Blood (Northeast Laboratory Service, P1100), Pseudomonas Isolation Agar, or Sabouraud Dextrose Agar + 100 μg/ml gentamicin made according to the manufacturer’s directions. Plates were incubated for 24 to 48 h at 37°C then imaged.

In order to identify isolates that grew on plates, clinical isolates were struck to single colonies, grown as single-colony cultures in either liquid LB medium (bacteria), liquid YPD medium (*Candida* species), or liquid glucose minimal medium (*Aspergillus fumigatus*). Bacterial 16S rDNA (forward primer: GTGSTGCAYGGYTGTCGTCA; reverse primer: ACGTCRTCCMCACCTTCCTC) and fungal ITS1 region (forward primer: GTA AAA GTC GTA ACA AGG TTT C; reverse primer: GTT CAA AGA YTC GAT GAT TCA C) were PCR amplified, and the identity of each clinical isolate was determined after sequencing each amplicon using the Applied Biosystems 3730 DNA analyzer (ThermoFisher Scientific).

### Statistics

Significance of correlation to determine reproducibility and linearity of microbial detection by NanoString was calculated using linear regression in Graphpad Prism 5. Correlation between NanoString and HiSeq sequencing were using R to determine the Pearson *r* correlation coefficient and its p-value for each subject across selected taxa and for the selected taxa across all subjects using the “cor.test” function in the “stats” package (v3.3.0) with default settings. Significance of correlation between *Candida* spp. colony numbers and NanoString counts and the significance of differences were calculated using linear regression or the Mann-Whitney test in GraphPad Prism 5.

## Acknowledgements

Research reported in this publication was also supported by grants from the National Institutes of Health to D.A.H. (R01 GM108492 to DAH). Support from the Cystic Fibrosis Foundation Research Development Program (STANTO011R0) as a pilot grants to J.D.S., A.H.G. and D.A.H.. The Dartmouth Lung Biology Center and CF Translational Research Core was supported by an Institutional Development Award (IDeA) from the National Institute of General Medical Sciences of the National Institutes of Health under grant number P30GM106394 and by the CFF RDP (CFRDP STANTO11R0). A.H.G. was supported by the CF Foundation Clinical Research Scholars Program under award number GIFFOR17Y5 and the Dartmouth Clinical and Translational Science Institute (SYNERGY) under award number KL2TR001088 (Green, P.I.) from the National Center for Advancing Translational Sciences (NCATS) of the National Institutes of Health (NIH). The content is solely the responsibility of the authors and does not necessarily represent the official views of the NIH.

---

**Fig. S1.**
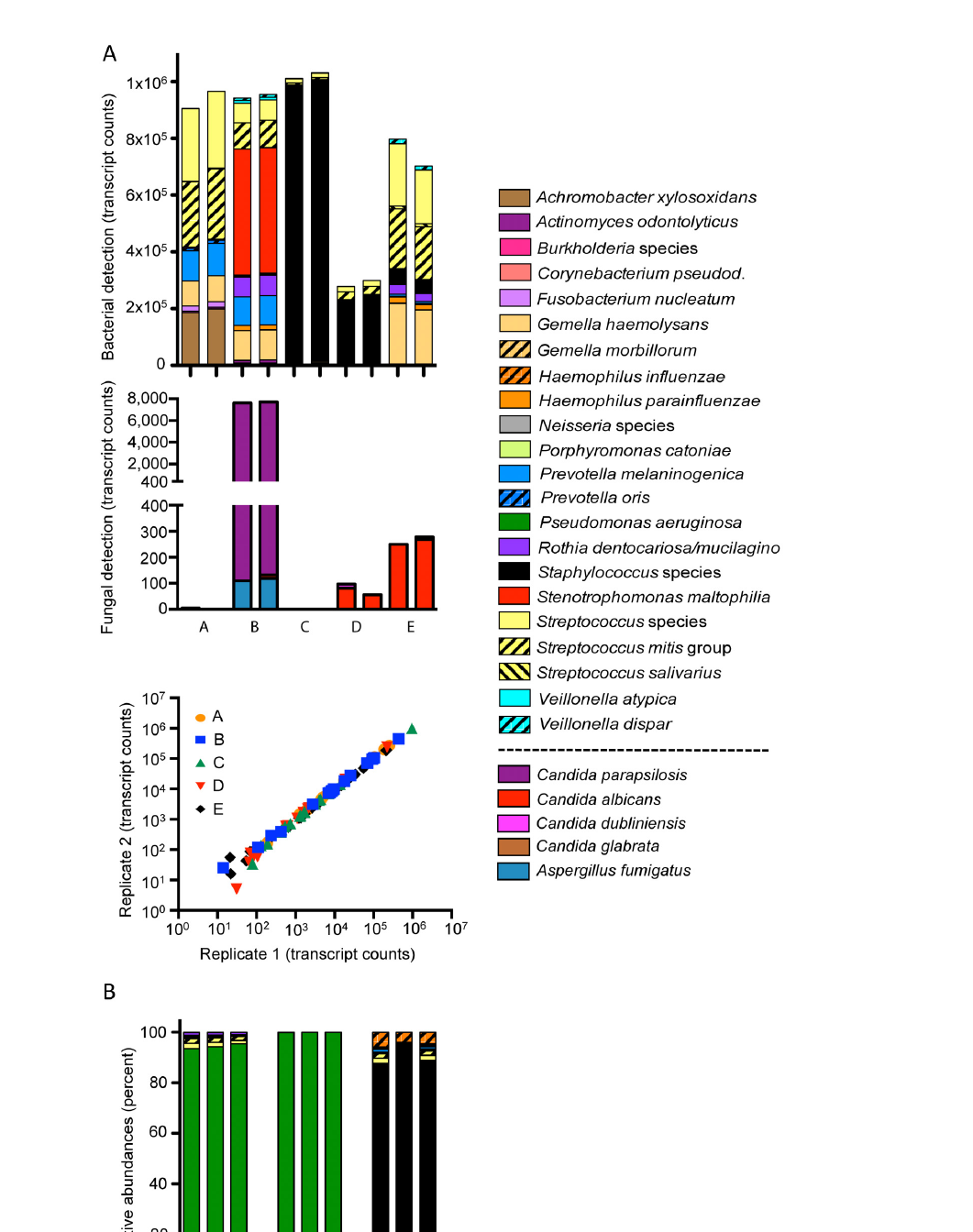
Reproducibility of NanoString analysis of sputum. **(A)** Microbial profiles in CF sputum RNA (sputum samples A-E) analyzed in duplicate by NanoString analysis are shown. Bacteria and fungi are presented separately. Correlation of count numbers by NanoString between duplicates for samples (A-E). **(B)** Microbial community profiles presented as percent abundance in three replicate RNAs extracted in parallel from three separate sputum samples (F, G, and H). The legend is the same as in A.

**Fig. S2.**
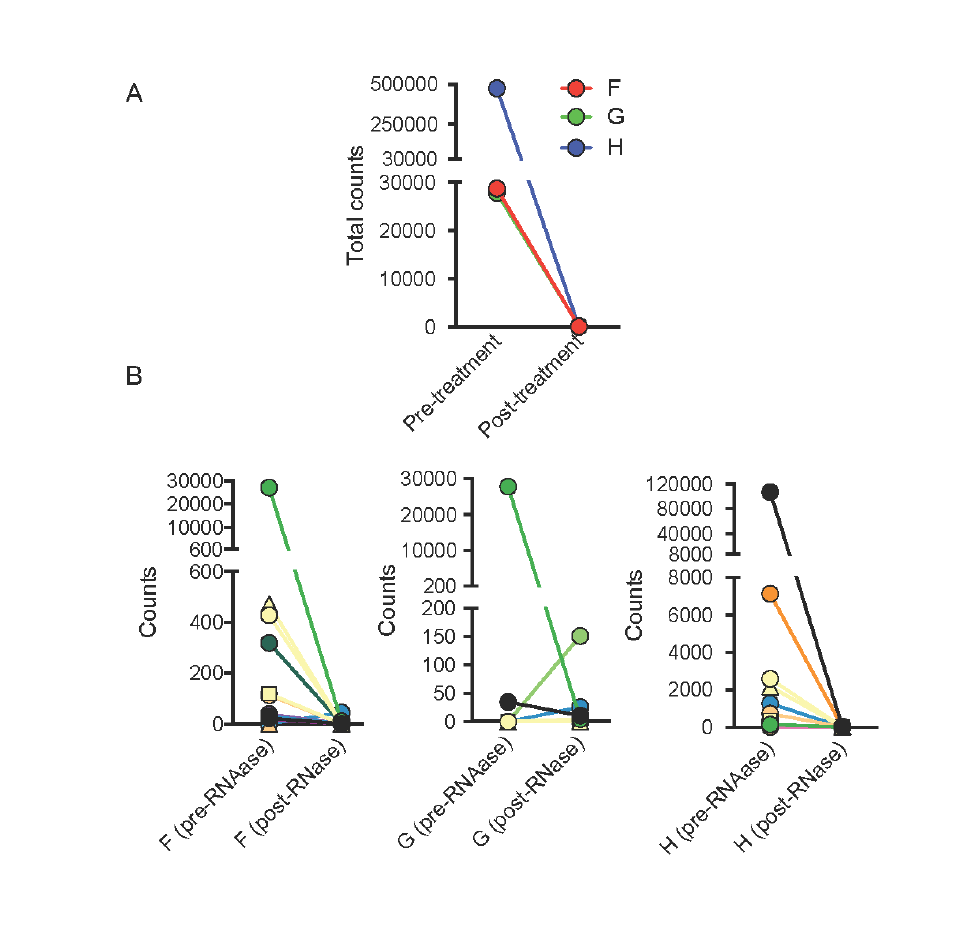
RNAse treatment of extracted RNA depletes detection of patient sample microbial communities by NanoString. Microbial communities were detected from total RNA of three sputum samples (F, G, and H) pre-and post-RNAse treatment with NanoString probes. **(A)** Total detection (sum of counts from all probe sets) from pre-and post-RNAse-treated samples. **(B)** Probe-specific transcript detection pre-and post-RNAse-treatment. Different species/genera detected are indicated by the different colors and symbols. The data values are presented in **Table S3B**.

**Fig. S3.**
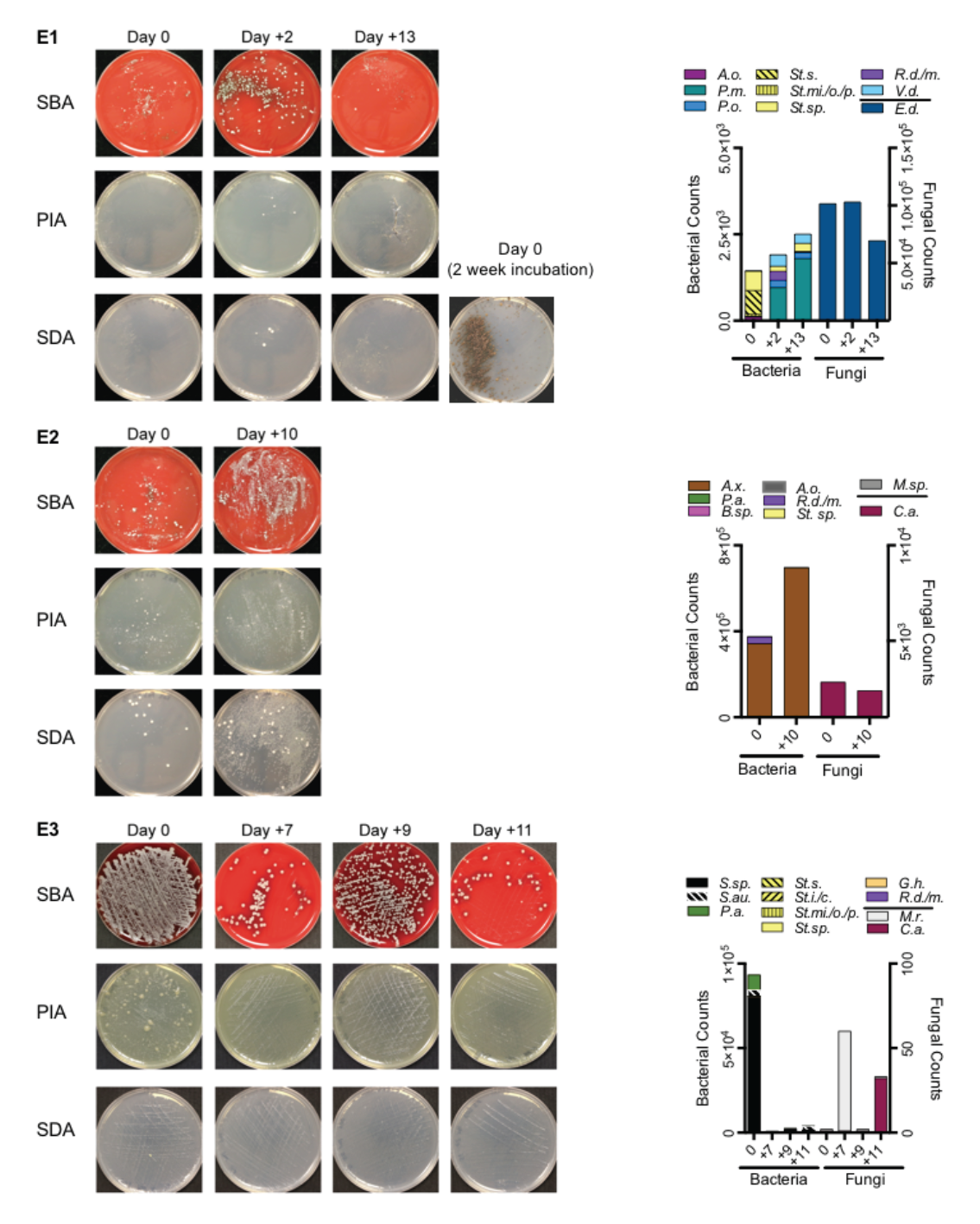
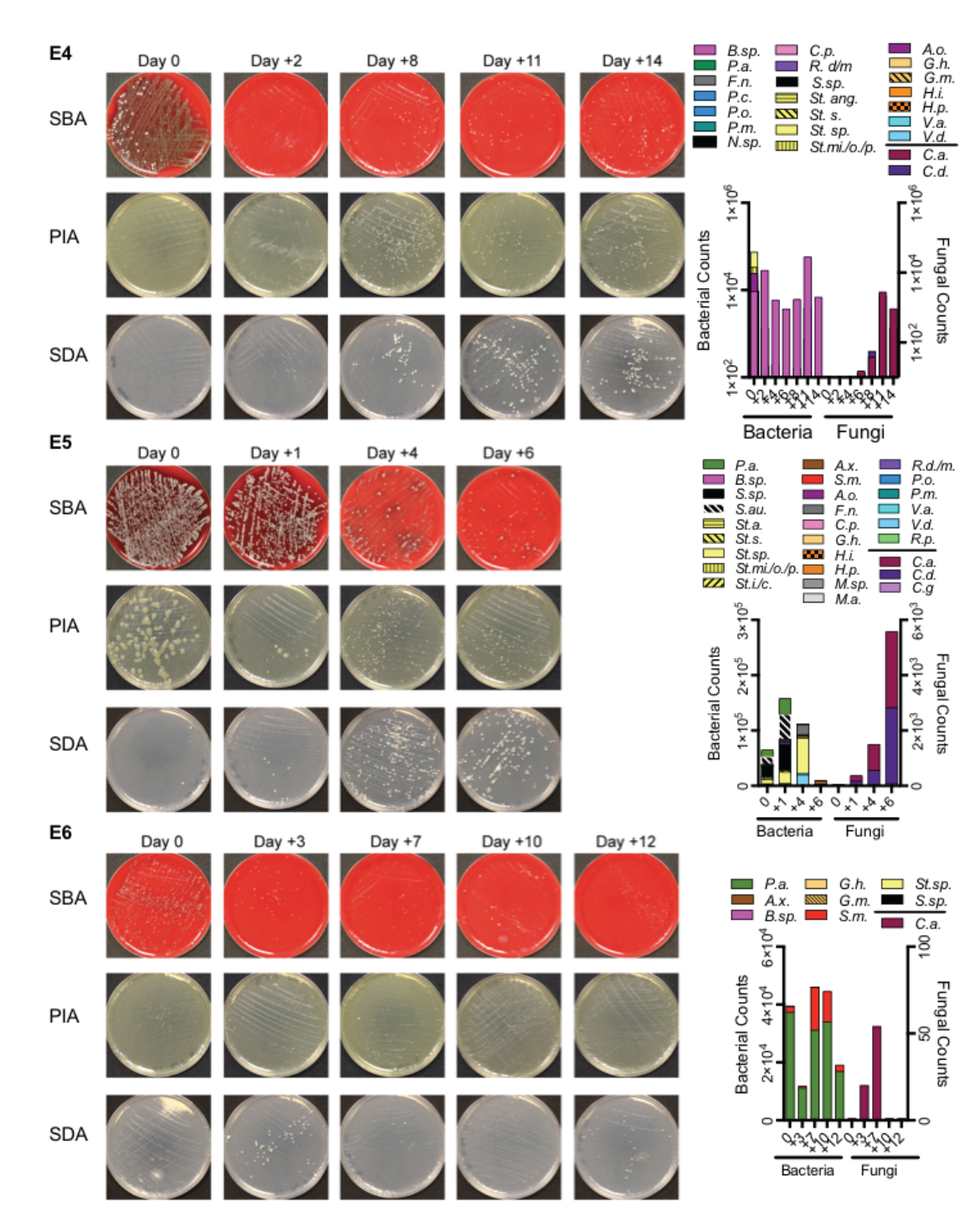
Culture analysis and NanoString analysis of sputum series from six hospitalized patients. Culture of E1-E6 sputum series on sheep blood agar, PIA, and Sabouraud dextrose agar plates. Plates were incubated for at least 48 h before imaging. Samples were collected on Day 0 (date of admission for treatment of exacerbation) and the days post treatment is shown for subsequent samples. For each series, RNA was analyzed by NanoString and data above background are shown for the bacterial and fungus targeting probes included in the code set. Exacerbation series E1 showed high levels of *Exophiala dermatitidis* in plate cultures only after extended incubation for all days and the plate from Day 0 is shown as an example. The clinical culture data for the day 0 samples are shown in **Table 2**. The NanoString data in the graphs for each sample are the same data as shown in Fig. 3, and are presented here for ease of comparison.

**Table S1**. NanoString probe sets designed for this study.

**Table S2**. Strains used in this study.

**Table S3.** All NanoString counts for samples.

**Table S4**. Illumina Reads and Illumina vs NanoString.

